# Colonization of the murine oropharynx by *Streptococcus pyogenes* is governed by the Rgg2/3 quorum sensing system

**DOI:** 10.1101/2020.06.30.179903

**Authors:** Artemis Gogos, Michael J. Federle

## Abstract

*Streptococcus pyogenes* is a human-restricted pathogen most often found in the human nasopharynx. Multiple bacterial factors are known to contribute to persistent colonization of this niche, and many are important in mucosal immunity and vaccine development. In this work, mice were infected intranasally with transcriptional regulator mutants of the Rgg2/3 quorum sensing (QS) system—a peptide-based signaling system conserved in sequenced isolates of *S. pyogenes*. Deletion of the QS system’s transcriptional activator (Δ*rgg2*) dramatically diminished the percentage of colonized mice while deletion of the transcriptional repressor (Δ*rgg3*) increased the percentage of colonized mice compared to wild type. Stimulation of the QS system using synthetic pheromones prior to inoculation did not significantly increase the percentage of animals colonized, indicating that QS-dependent colonization is responsive to the intrinsic conditions within the host upper respiratory tract. Bacterial RNA extracted directly from oropharyngeal swabs and evaluated by quantitative RT-PCR subsequently confirmed QS upregulation within one hour of inoculation. In the nasal-associated lymphoid tissue (NALT), a muted inflammatory response to the Δ*rgg2* bacteria suggests that their rapid elimination failed to elicit the previously characterized response to intranasal inoculation of GAS. This work identifies a new transcriptional regulatory system governing the ability of *S. pyogenes* to colonize the nasopharynx and provides knowledge that could help lead to decolonization therapeutics.

**Author Summary:** *Streptococcus pyogenes* is responsible for a wide spectrum of diseases ranging from common pharyngitis to infrequent invasive infections like necrotizing fasciitis. The ability of this microorganism to persist in the human oropharynx predisposes colonized individuals to a variety of superficial and invasive diseases which lead to significant morbidities and mortality. Identification of the regulatory systems that augment the bacteria’s ability to colonize the oropharynx provides potential targets against which molecular therapeutics can be designed. Here we show that the Rgg2/3 quorum sensing system, an interbacterial communication system, governs the ability of *S. pyogenes* to colonize the murine oropharynx. Disruption of the system’s transcriptional activator reduced colonization dramatically, eliminated the transcription of two sets of genes known to be activated by the Rgg2/3 system, and tempered the innate immune response seen when *S. pyogenes* is intranasally infected into the mouse.

## Introduction

*Streptococcus pyogenes*, or Group A Strep (GAS), is a human-restricted pathogen able to colonize and adapt to numerous tissues of the body. The oronasopharynx is regarded as the most common site of colonization by GAS, which can lead to acute infections such as streptococcal pharyngitis. GAS is responsible for 600 million annual cases of symptomatic pharyngitis, and a meta-analysis estimates that 8-40% of children are asymptomatically colonized [1]. It is crucial to identify factors governing the persistence of the bacteria in this niche to aid in the development of anti-virulence treatments and vaccines. Work by others has already identified some host and bacterial factors that contribute to GAS colonization. On the bacterial side, colonization is aided by multiple virulence factors, many of which are controlled by the CovRS regulatory system and *mga* [2–6]. For example, the capsule, which is under control of CovRS, has been shown to be critical for pharyngeal colonization in both primate and murine models [7, 8]. Animal models have shown that the host also plays a role in mediating GAS colonization and preventing bacteria from disseminating systemically. During pharyngeal infection, an influx of neutrophils and increased proliferation of CD4+ T cells are seen in the nasal associated lymphoid tissue (NALT) of the mouse, which is analogous to human tonsils [9]. Further review of known bacterial and host factors important for GAS oropharyngeal colonization have been discussed recently [10].

In many other bacterial species, the ability to colonize a host is aided by cell-to-cell communication systems, also commonly known as quorum sensing (QS) [11] [12–14] [15–18]. These systems function by way of sensing bacterially produced, extracellular small molecules or peptides, referred to as autoinducers or pheromones, that disseminate information to coordinate gene expression and behaviors among members of a bacterial population. Although the transcriptional targets of QS systems vary significantly between bacteria, many distantly related species utilize these systems to adapt to specific niches in the host. In this work, we sought to build upon this knowledge and determine whether a quorum sensing system in GAS, the Rgg2/3 system, is involved in colonization of the nasopharynx. The Rgg2/3 system in GAS consists of two transcriptional regulators, an activator (Rgg2) and a repressor (Rgg3), that are modulated by two, nearly identical short hydrophobic peptide signals (SHPs) which interact and bind with the two Rgg regulators (**Fig. 1**) [19, 20]. At low concentrations of peptide signal, Rgg3 predominates as a negative transcriptional regulator [19]. When pheromones accumulate to levels that result in receptor binding, they bind to and evoke conformational changes in both Rgg proteins, ceasing repression by Rgg3 while inducing Rgg2 to activate transcription. The primary targets of activation are two operons that include the genes encoding the SHPs, and thus, the net effect of SHP binding to Rgg2 and Rgg3 is robust transcriptional induction resulting in a positive-feedback regulatory loop made possible by further pheromone production. An additional regulatory element of this circuit is the PepO peptidase, a CovRS-regulated gene that functions to digest the peptide pheromones and turn off the transcription of Rgg2/3 targets [21].

**Figure 1.**
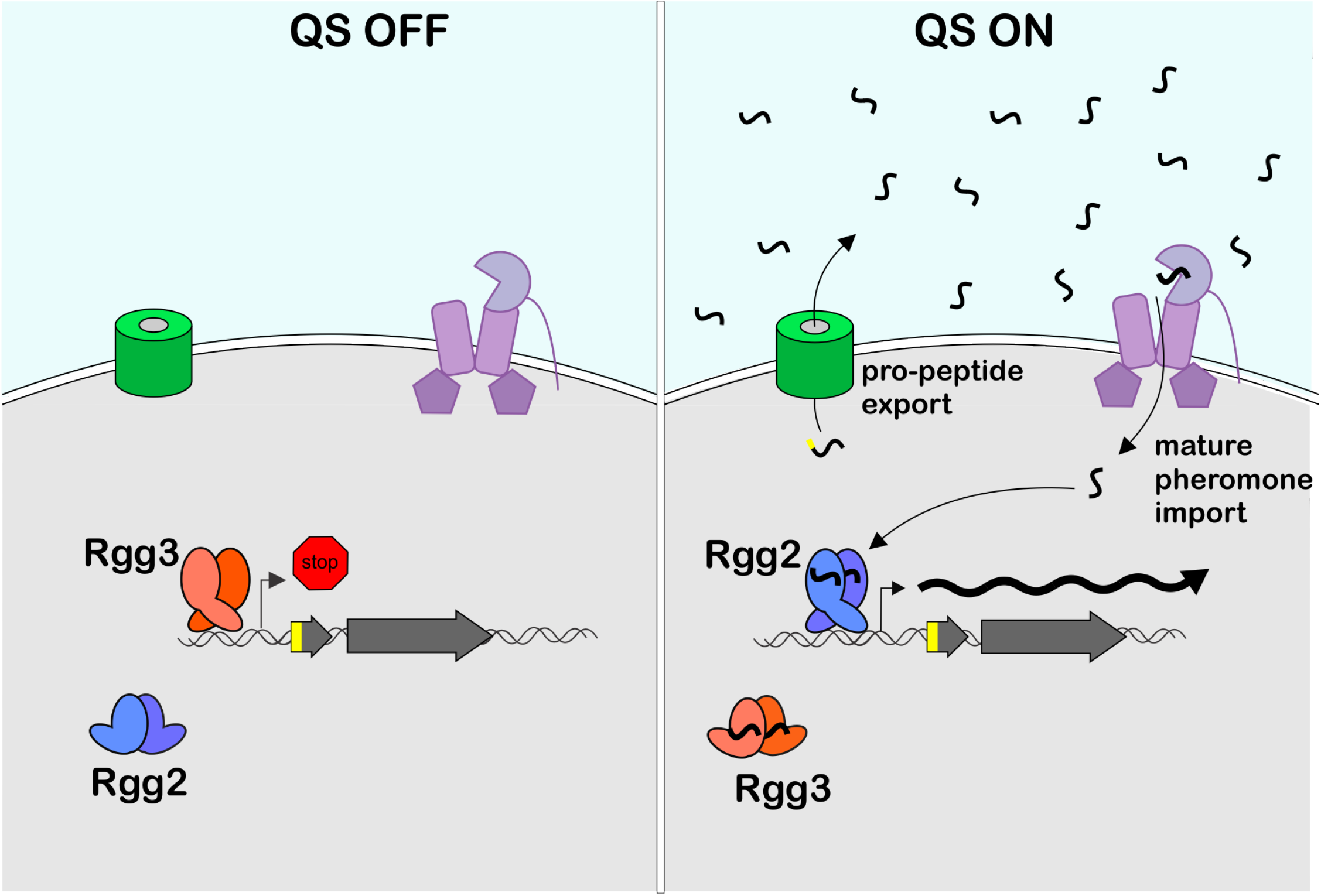
The Rgg2/3 Quorum Sensing Circuit. The molecular architecture of the Rgg2/3 quorum sensing system in *Streptococcus pyogenes* consists of short hydrophobic peptide (SHP) pheromones that are processed, exported and reimported into the cytoplasm where they can bind to two transcriptional regulators: the activator, Rgg2, and the repressor, Rgg3. Deleting Rgg3 prevents repression and moderately increases expression of target genes, whereas deletion of Rgg2 disables transcription of target promoters.

Initial activation of the Rgg2/3 system can be stimulated by certain environmental conditions, such as low levels of iron and manganese, or the presence of mannose as the primary carbon source [22]. There is evidence for each of these environmental conditions to be present in the pharyngeal mucosa. Metals are generally considered to be restricted by the host and sparingly available to bacteria [23], although to our knowledge, exact mucosal measurements of iron and manganese have not been studied. However, it is known that mannose is present in the Man_3_GlcNAc_2_ core of N-glycan decorations on the surface of nasopharyngeal epithelial cells [24]. *S. pneumoniae* is able to break down these decorations using a number of mannosidases [25], of which homologs exist in GAS [26]. Additionally, it has been confirmed that GAS pharyngeal isolates from cynomologous macaques upregulate mannose catabolism genes, indicating that mannose is likely available to GAS in the pharynx [27]. Due to these environmental conditions, we have previously hypothesized that the Rgg2/3 regulatory system may be playing an important role in host-bacterial interactions [21, 22, 28]. In this work, we present evidence showing that the Rgg2/3 QS system governs pharyngeal colonization in the mouse via activation of its downstream gene targets.

## Results

### Colonization of the murine oropharynx by *Streptococcus pyogenes* requires an intact Rgg2/3 QS system

Prior studies have shown that serotype M3 strain MGAS315 colonizes mice well and for relatively long periods of time [Flores et al.], making it an ideal strain to test colonization phenotypes in the mouse. MGAS315 is a *covS* mutant [29–31] and displays higher expression of virulence factors, including higher PepO levels. As PepO degrades SHP pheromones, we reasoned that SHP-dependent signaling activity would remain at low levels until the system can be activated by a favorable environment [21].

To confirm QS activity in MGAS315, RNA levels of the *stcA* gene—one of the main targets of the Rgg2/3 system—were used as a reporter of QS activity and measured by quantitative RT-PCR (qRT-PCR) from wild type and isogenic strains containing deletions of either *rgg2* or *rgg3*. Cultures were grown in a chemically defined medium (CDM) to determine the baseline activity of the quorum sensing system prior to inoculation of the mouse. Without the addition of synthetic pheromone, all three strains displayed similar, low levels of expression of the *stcA* gene (**Fig 2A**). However, addition of synthetic pheromone (SHP-C8) to cultures led to rapid and robust induction of the *stcA* gene in wild type and Δ*rgg3*, consistent with the model that Rgg3 is not required for activation. Conversely, the Δ*rgg2* mutant was not able to turn on QS targets, regardless of the addition of pheromone, again in line with the idea that Rgg2 is required for activation.

**Figure 2.**
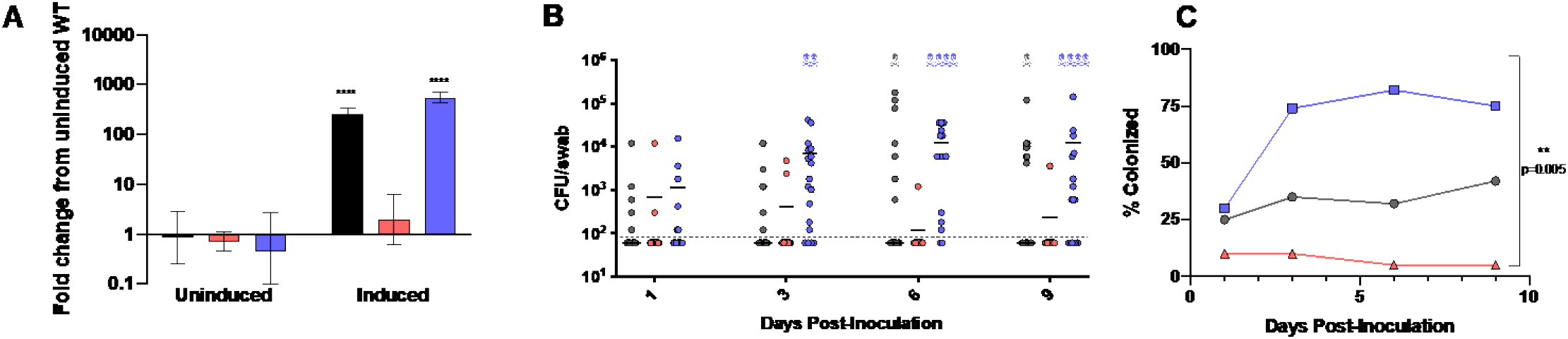
Colonization of the murine oropharynx by *Streptococcus pyogenes* requires an intact Rgg2/3 quorum sensing system. The MGAS315 serotype of GAS was analyzed for QS activity in culture and for colonization levels in the mouse. (A) Relative transcript levels of QS-target gene *stcA* in cultures of WT, Δ*rgg2*, and Δ*rgg3* strains grown in CDM broth without (uninduced) or with (induced) the addition of 100 nM SHP-C8 pheromone. Bars are the mean fold differences compared to uninduced WT for three biological replicates. Asterisks indicate significance of multiple t-tests comparing each sample to uninduced WT, α≤0.05. (B) Three-week old CD1 mice were inoculated intranasally with with 10^7^ CFU of uninduced bacteria. CFU counts obtained from throat swabs of each mouse are plotted for each day of sampling. Skull and crossbones denote the number of animals that displayed humane endpoint signatures requiring euthanasia (signs of distress indicating invasive infection) and are shown in an accumulating manner. Twenty mice were utilized for each bacterial strain; bars indicate the median CFU/swab value and the detection limit was 60 CFU/swab. Using the Friedman test, a nonparametric matched ANOVA, the three strains were determined to have statistically different median CFU counts over the course of the experiment, p=0.028. (C) The percentage of mice colonized during the experiment shown in Fig 2B were compared between the three strains over time. Deceased mice were not included during the calculation of percent colonized mice. A Friedman test was again run to compare the colonization percentages between the three strains, and the colonization rates of each strain were found to be statistically significant from one another, p=0.0046.

We then examined the importance of an intact Rgg2/3 QS system during murine oropharyngeal colonization by inoculating CD1 mice intranasally with uninduced wild type, Δ*rgg2*, or Δ*rgg3* bacteria and monitored the colonization load over 9 days (**Fig 2B, 2C**). Wild type bacteria were able to colonize the upper respiratory tract for the entire experimental course. Bacteria with a deletion of the system’s transcriptional activator, *rgg2*, were unable to colonize the oropharynx in nearly all inoculated mice. In the few Δ*rgg2*-infected mice with recoverable bacteria, CFU counts of Δ*rgg2* were far lower than seen in mice inoculated with wild type. Curiously, Δ*rgg3* strains were found to colonize higher proportions of mice with higher numbers of CFUs when compared to wild type. Of the twenty mice infected with the Δ*rgg3* strain, four mice displayed experimental endpoint characteristics of invasive disease on the third and sixth day and were euthanized. As CFU counts were among the highest and in the largest proportion of mice for the Δ*rgg3* strain, it stands to reason the probability for invasive infection were highest in these mice.

Although the Rgg2/3 system is conserved in all sequenced strains of GAS, signaling dynamics and efficiency is not equivalent among strains tested [19, 21, 22].Therefore, two other GAS serotypes were tested to evaluate the contribution of these regulators in the nasopharyngeal colonization model. Strains NZ131 (serotype M49) and HSC5 (M14), and their isogenic *rgg2* and *rgg3* mutants, were inoculated in mice and monitored periodically by throat swab and CFU counting. Although these serotypes were less informative than MGAS315 due to differences in burden and length of bacterial infection, results indicated that the Δ*rgg2* derivatives were attenuated in their ability to colonize and confirmed that the system governs the same responses in other clinical isolates of *S. pyogenes* (**Fig S1**). Furthermore, these additional strains confirmed that the Δ*rgg3* mutants colonized better than WT.

### Pre-activation of the QS system before inoculation does not affect colonization outcome

We next tested whether wild type bacteria would colonize to levels similar to the Δ*rgg3* mutant if they were pre-stimulated with pheromone prior to inoculation. We reasoned that pre-stimulation might enhance the conditioning of bacteria for pharyngeal colonization, producing proteins that would alter the bacterial surface or physiology. We repeated the colonization experiments as described above, with the addition of a cohort of mice inoculated with QS-activated cultures that had been incubated with SHP-C8 pheromone for one hour prior to inoculation. When stimulating MGAS315 cultures for this amount of time, the QS target genes are induced by nearly 1000-fold (**Fig. 2A**). Following nasal inoculation, throat swab monitoring over a nine-day period showed no significant change in colonization between pre-stimulated and unstimulated cultures (**Fig. 3**), indicating that the status of QS activity prior to inoculation did not affect the establishment of colonization or mortality.

**Figure 3.**
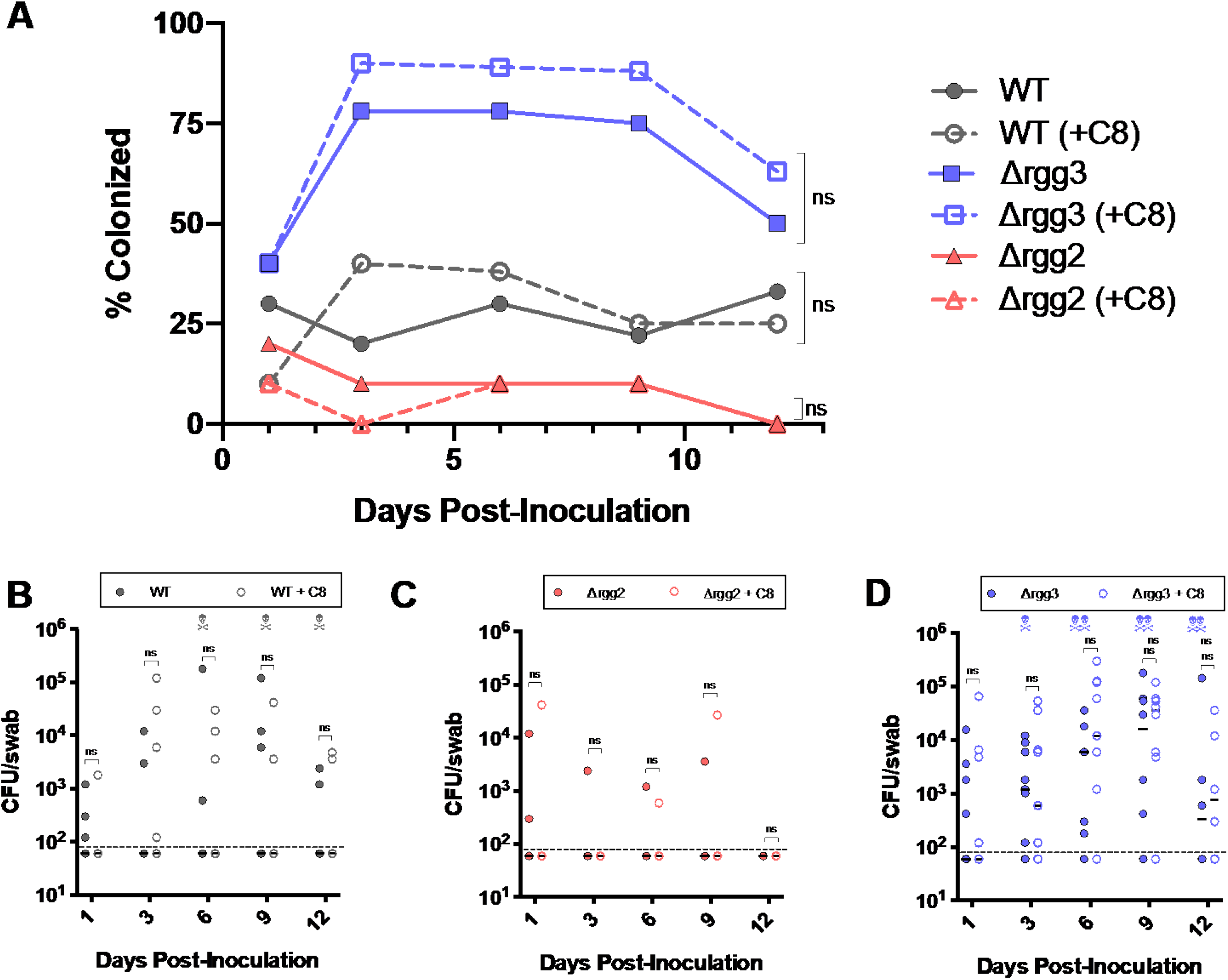
Pre-activation of inoculated bacteria does not alter the outcome of colonization. Ten mice per condition were inoculated with MGAS315 bacterial strains grown in CDM without or with (+C8) the addition of 100 nM SHP-C8 pheromone in the medium, or 1μl DMSO for the control. The cells were resuspended in PBS with 100 nM SHP-C8 pheromone. Upon inoculation, no additional pheromone was administered to the mice. Throat swabs were taken for 12 days. (A) The percentages of mice colonized with or without the addition of SHP-C8 were compared using a Wilcoxon matched-pairs signed rank test (α≤0.05). (B-D) Colony CFU counts for WT, Δ*rgg2*, and Δ*rgg3* were also graphically and statistically compared between the control and pheromone-treated groups. The colony counts on each day were compared using multiple t-tests (α≤0.05).

### Intranasal bacterial inoculation leads to the induction of the Rgg 2/3 QS system

Given the above evidence that *in vivo* conditions impact quorum sensing, we were eager to measure the activity of the QS system of bacteria within the host. As an initial means to visualize QS activity in living animals, we inoculated mice intranasally with unstimulated MGAS315 wild type, Δ*rgg2*, or Δ*rgg3* strains containing a multicopy P_*shp3*_*-luxAB* reporter before undergoing anesthesia for immobilization. We then obtained imaging at 15- and 60-minutes post-infection using an In Vivo Imaging System (IVIS). Imaging showed that luciferase activity increased over time in mice inoculated with wild type and Δ*rgg3* bacteria (**Fig. 4A, 4C**), but those inoculated with the Δ*rgg2* mutant displayed a lower luciferase intensity and the amount did not change over time (**Fig. 4B**).

**Figure 4.**
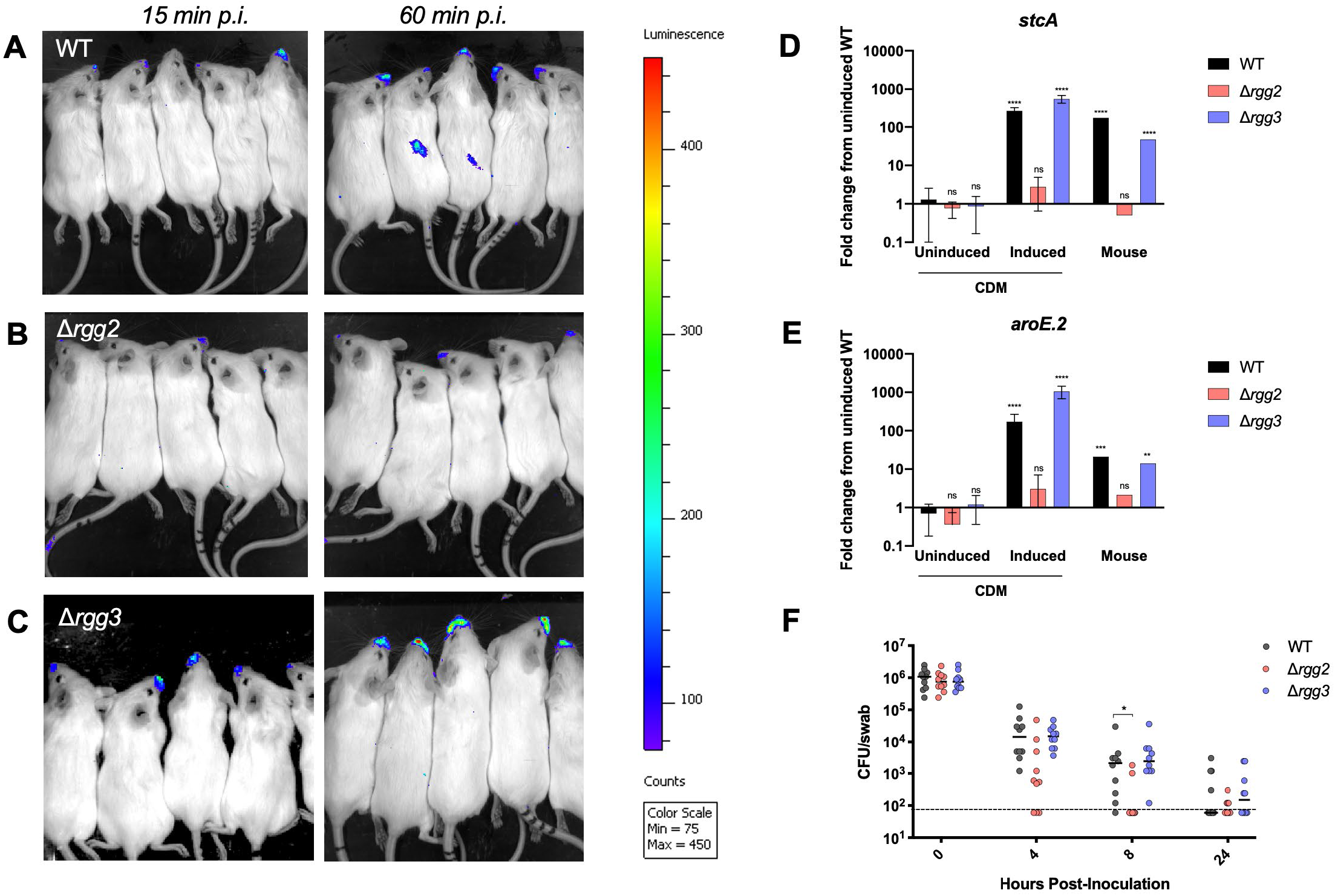
The Rgg2/3 regulon is induced upon intranasal inoculation into the mouse. (A-C) Mice intranasally inoculated with MGAS315 bacteria containing the pJC229 QS luciferase reporter were imaged 15- and 60-minutes post-infection using an *in vivo* Imaging System (IVIS). Transcriptional expression of the *stcA* and *aroE.2* (E) quorum sensing targets in CDM without (uninduced) or with (induced) the addition of 100nM SHP-C8 peptide pheromone, as well as from RNA extracted from the pooled throat swabs (mouse) of 10 mice swabbed one hour after infection. The mean fold change and error bars are shown in the CDM conditions for three biological replicates. The mouse samples were pooled from 10 throat swabs and the bars represent the average of three technical replicates, thus no error bars are shown. The Holm-Sidak method was used to determine which conditions produced transcript levels significantly different from wild-type in uninduced CDM. ****p<0.0001, ***p<0.001, **p<0.01 (F) Bacterial counts from throat swabs taken 0-24 hours post-inoculation from mice inoculated with MGAS315 Δ*rgg3* or MGAS315 Δ*rgg2*. The median Δrgg2 and Δrgg3 colony counts were compared to the WT counts using a Wilcoxon test (α≤0.05). The Δrgg2 colony counts were statistically different from WT at 8 hours (p=0.0273) post-inoculation.

To validate imaging results and to obtain a more precise quantification of transcription of QS-regulated genes during early stages of colonization, we isolated bacterial RNA directly from mouse throat swabs and conducted qRT-PCR. Three groups of ten mice were inoculated with either wild type, Δ*rgg2*, or Δ*rgg3*. One hour after inoculation, swabs of each group were combined and extracted with TRIzol to isolate nucleic acid. To evaluate QS activation, we assessed transcript levels (normalized to the housekeeping gene *gyrA*) of two QS-targeted genes, *stcA* and *aroE.2,* from swab samples and compared them to transcript levels from the uninfected inoculum (**Fig. 4D, 4E**). As expected, both *stcA* and *aroE.2* were highly upregulated in wild type and Δ*rgg3* strains that were induced with pheromone in vitro. In bacteria collected from mice, we observed a similar pattern, with wild type and Δ*rgg3* showing high upregulation of *stcA* and moderate upregulation of *aroE.2*. The Δ*rgg2* strain displayed no induction of *stcA or aroE.2* expression and their levels were similar to samples collected from in vitro cultures, indicating that *rgg2* is responsible for the increased transcription of *stcA* and *aroE.2* in the mouse. Although in vivo expression of *stcA* and *aroE.2* in wild type and Δ*rgg3* was lower than in induced in vitro cultures, it was nonetheless highly upregulated compared to the unstimulated inoculum.

As the IVIS and qRT-PCR data suggested that the Rgg2/3 system is important within the first hour of colonization, we examined the rate of bacterial clearance during the first 24 hours post-inoculation. We performed throat swabs at 0, 4, 8, and 24 hours post-inoculation and showed that even by four hours most bacteria, especially the Δ*rgg2* cells, were already being eliminated from the oropharynx (**Fig. 4F**). The rapid clearance suggests that elimination of bacteria is by a mechanism consistent with humoral components such as antimicrobial peptides (AMPs) and lysozyme, or primary immune mechanisms such mucociliary clearance and swallowing. Although WT and Δ*rgg3* bacteria were better adapted at surviving this reduction, they too were reduced by several orders of magnitude over 24 hours, congruent with data from **Fig. 2B** that shows lower bacterial burden on day 1.

### Quorum sensing non-compliance in the bacterial population does not alter colonization patterns

Quorum sensing systems fundamentally exist to provide bacteria a mechanism to coordinate gene activity across a population. We reasoned that for GAS, the host immune system provides the primary obstacle to establishing colonization, hence a coordinated response by bacteria against host immunity would benefit the population, possibly through a mechanism of coordinated immune avoidance or coordinated assault on immune function. We expected that bacteria that do not (or cannot, in the case of Δ*rgg2* bacteria) comply with quorum activities would be a detriment to the population, either because immune responses would become activated or because non-participating bacteria would not contribute to impairing immune activities, especially if they are a substantial component of the population. Given that Δ*rgg2* bacteria are unable to participate in the coordination of gene responses, we predicted that inoculating mice with a mixture of Δ*rgg2* and Δ*rgg3* bacteria would have one of two outcomes: Either the non-compliant Δ*rgg2* bacteria would have a negative impact on the suitability of the niche (possibly by causing immune activation), leading to decreased colonization by both strain types; or Δ*rgg3* cells would compensate for the deficiencies of Δ*rgg2* (possibly through trans-complementation of secreted factors regulated by the quorum sensing system), thus increasing the ability of individual Δ*rgg2* bacteria to colonize. To test these scenarios, mice were co-inoculated with both Δ*rgg2* and Δ*rgg3* mutants at a 1:1 ratio and pharyngeal swabs were taken over a nine-day period. Surprisingly, neither predicted outcome was observed, and instead, bacterial colonization by each mutant mirrored the results of when strains were inoculated individually. By the first day, mice were nearly exclusively colonized by Δ*rgg3* bacteria, whose numbers continued to expand over the first week; Δ*rgg2* bacteria were cleared by day 3 (**Fig. 5A**). Co-inoculations were also carried out for mixtures of wild type with Δ*rgg2*, which also led to the elimination of *Δrgg2* bacteria. This combination saw wild type bacteria colonize a moderately higher percentage of the mice, and to higher CFU levels, than the single infection experiment, although these results were not statistically significant (**Fig. 5B**). Co-inoculation of wild type with Δ*rgg3* bacteria also showed no statistically significant differences in colonization patterns from those observed in the single infections (**Fig. 5C**). Thus, results from these experiments point to a more complex situation occurring in the host environment. One possible confounding factor is that Δ*rgg2* bacteria are eliminated more rapidly than wild type or Δ*rgg3,* and this rapid removal may not provoke an immune response to an extent where there is a negative impact on colonization by wild type or Δ*rgg3* bacteria.

**Figure 5.**
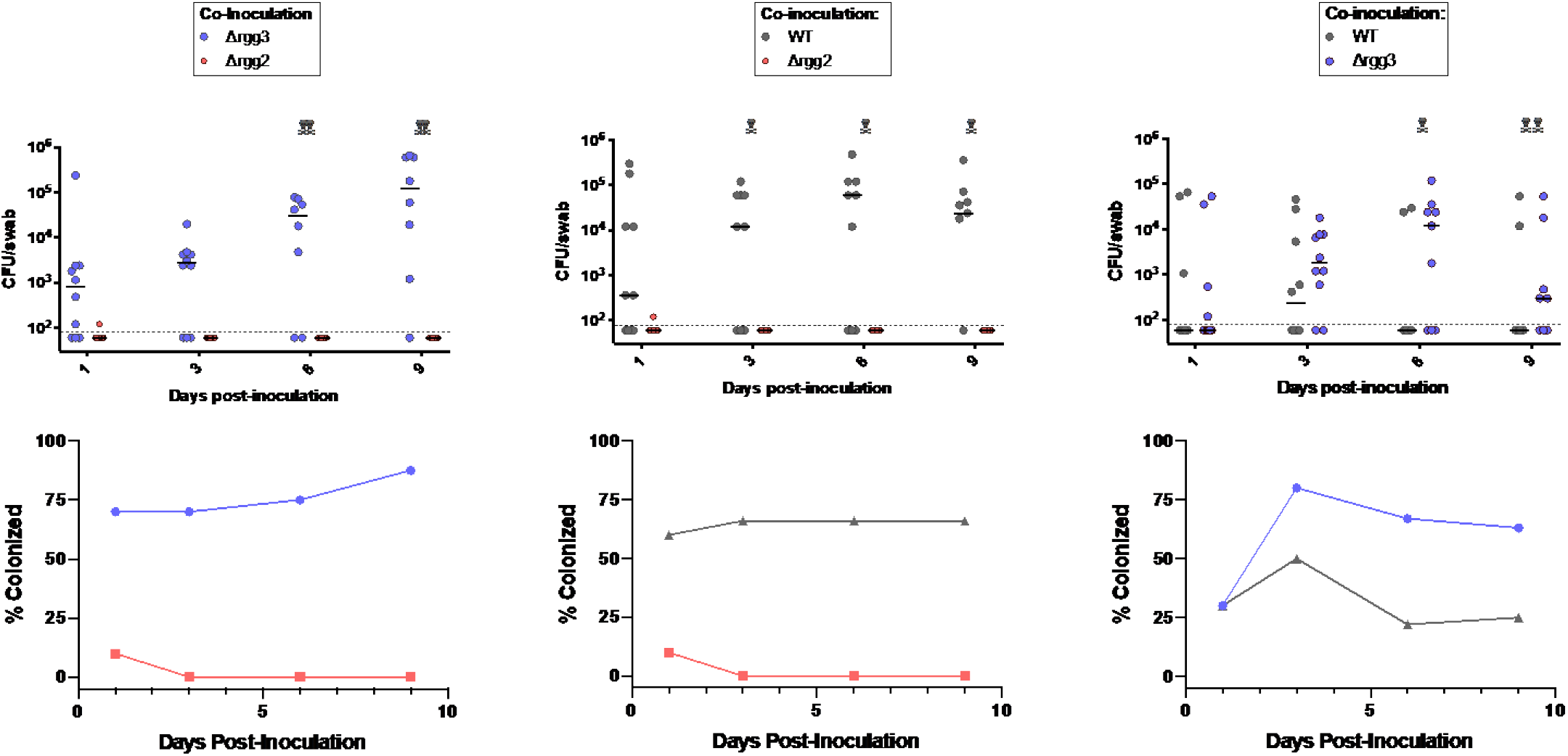
Co-inoculation of mice with QS mutants. 10^7^ CFU of each indicated MGAS315-derived strain was co-inoculated intranasally into the mouse. Ten mice per condition were sampled by throat swabbing for CFU counts. Strains were differentiated by growth on selective medium (WT: BD Strep Selective Agar, Δ*rgg2*: THY + 100μg/ml kanamycin, Δ*rgg3* : THY + 3μg/ml chloramphenicol). (A) Co-inoculation of MGAS315 Δ*rgg2* and Δ*rgg3*. (B) Co-inoculation of MGAS315 WT and Δ*rgg2*. (C) Co-inoculation of MGAS315 WT and Δ*rgg3*. Bars denote the median CFU value; skull and crossbones denote a deceased mouse and number of symbols indicate total accumulating dead mice over time. Limit of detection: 60 CFU/swab.

### The *rgg2* mutant fails to induce a pro-inflammatory response in the NALT

To further investigate the impact that Rgg2/3 QS has on host responses in this colonization model, we assessed bacterial loads and cytokine profiles from nasal-associated lymphoid tissue (NALT) of infected animals. For each of the three bacterial strains, we inoculated ten mice intranasally. At both 24 and 72 hours-post-infection, we swabbed the throats and surgically removed the NALT organ from each mouse. The NALT is a bi-lobed organ that was split into two equal halves; one half was homogenized and plated to obtain bacterial counts, and the other was processed to isolate secreted cytokines and chemokines, which were subjected to a bead-based multiplex assay for quantification.

Comparison of the CFU counts between the throat swabs and the NALT showed that all three bacterial strains were taken up into this lymphatic organ that drains the oropharynx and nasopharynx. At 24 hours post-infection, all three strains were present in the NALT of at least half of inoculated mice, although the Δ*rgg3* strain colonized in every instance and had on average 10-fold more CFU counts than either wild type or Δ*rgg2* (**Fig. 6B**). In the throat, bacterial counts of all strains were low at 24 hours (**Fig. 6A**), as seen in previous experiments. By day three, the Δ*rgg2* bacteria were no longer present in the NALT, and the NALT CFU counts mirrored the throat swab counts for all three strains (**Fig. 6A, 6B**). Cytokine response to the Δ*rgg2* cells, when compared to wild type, showed decreased expression of pro-inflammatory cytokines such as IL-1β and TNF-α (**Fig. 6D**) which have been previously described to be upregulated during GAS nasal colonization [32–34]. Likewise, the IFN-γ-inducible chemokine CXCL9, previously described to increase after GAS inoculation, was also significantly lower in the Δ*rgg2* inoculated mice when compared to mice with wild type GAS [33, 34]. Generally, it appeared that Δ*rgg2* bacteria failed to elicit cytokine responses and, taken together with the observed rapid clearance from pharyngeal and NALT tissues, may indicate that humoral effectors provide sufficient means to handle these bacteria without eliciting an inflammatory response.

**Figure 6.**
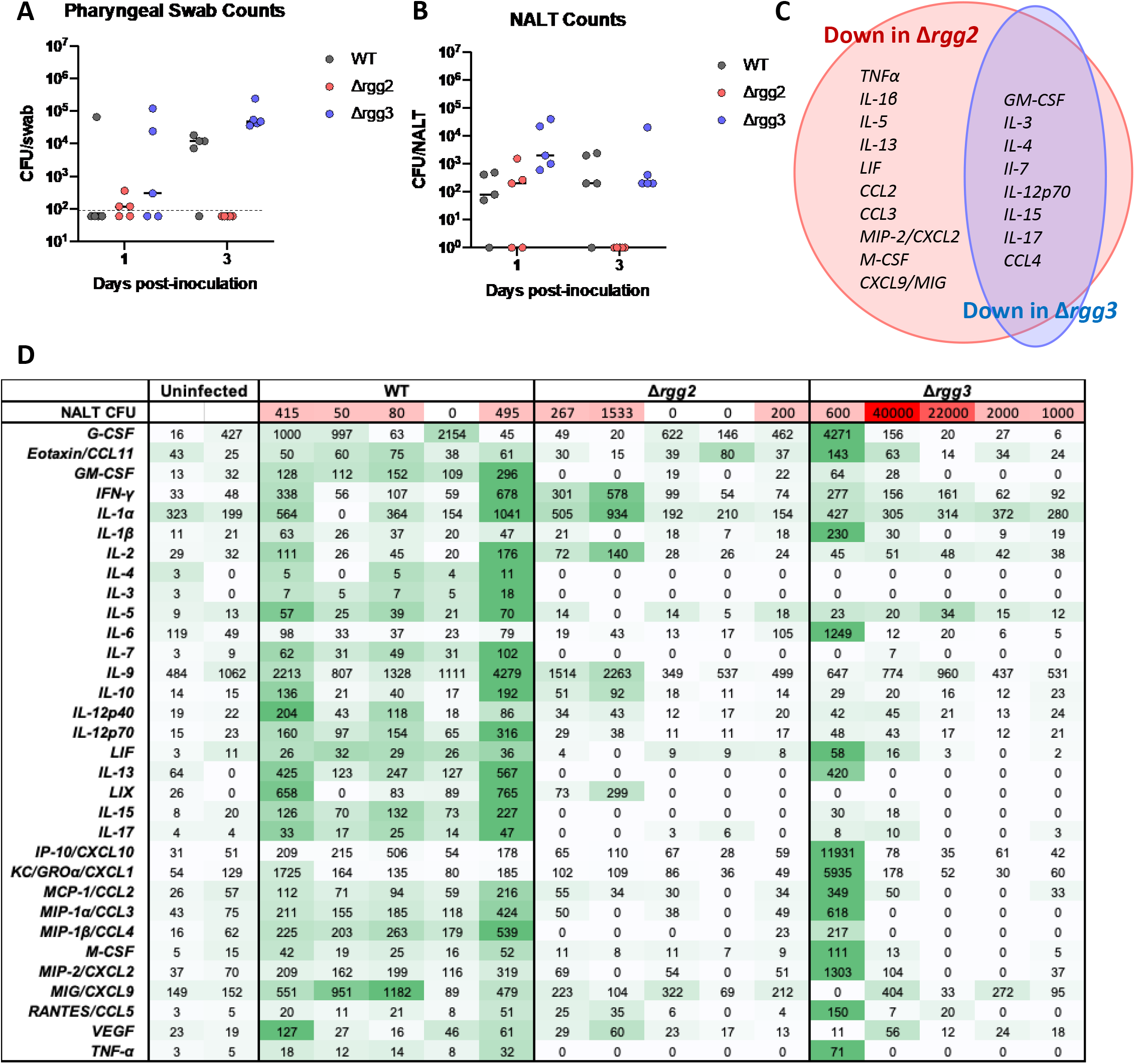
NALT response to intranasal inoculation of MGAS315 strains. 10^7^ CFU of each MGAS315 strain was inoculated intranasally into 10 mice. Five mice were swabbed and the NALT was dissected on day 1, the other five were swabbed and dissected on day 3. (A) Pharyngeal swab counts from the mice, limit of detection was 60 CFU. (B) CFU obtained from the NALT of each mouse on day 1 and day 3. (C) Venn diagram depicting the cytokines and chemokines that were significantly different from wild type for each *rgg* mutant strain. Data was obtained using a 32-plex cytokine array (Millipore) and t-tests were used to compare the protein levels between wild type and the mutants. (D) Raw values (pg/ml) obtained from each individual mouse 24 hours following inoculation for each of the 32 analyzed proteins. Zero values indicate measurements were below the limits of detection.

Immune responses to Δ*rgg3* bacteria were more difficult to interpret, as variability among mice was broader; this was primarily due to one mouse displaying unusually high cytokine and chemokine counts, even though all mice were colonized with high levels of bacteria. However, with exception of this mouse, cytokine and chemokine responses to Δ*rgg3* were generally lower compared to wild type, especially for a number of regulatory cytokines such as IL-3, IL-4, IL-7, and IL-17, which may help to explain the increased number of Δ*rgg3* cells that are able to survive both in the NALT and the throat.

## DISCUSSION

This work provides evidence that the Rgg2/3 quorum sensing system is required for oropharyngeal colonization by *S. pyogenes*. Congruent with our findings, a previous report showed that an insertional disruption in *rgg2* decreased the survival of mice in a model of intraperitoneal infection, indicating that this system may be broadly important for the host-pathogen interaction [35]. Recent evidence has shown that the RopB-SIP quorum sensing system and its cysteine protease product SpeB also contributes to GAS pharyngeal colonization in the nasopharynx [36, 37], indicating that cell-cell communication systems are important for this niche.

Curiously, deletion of the Rgg2/3 system’s repressor, *rgg3*, resulted in higher levels of colonization compared to wild type, although *in vivo* expression of the downstream target operons, *stcA and aroE.2*, was comparable between the WT and Δ*rgg3* strains (**Fig. 2, 4**). Perhaps transcription activation of targets diverges between the *rgg3* mutant and wild type at other times along the course of infections. The inability to alter the outcome of colonization by pre-stimulating wild type bacteria with synthetic pheromone (**Fig. 3**) indicates that the environment of the nasopharynx overrides any potential products that pre-conditioning produces during activation of the QS circuit. On a similar note, it was recently shown that although the competence quorum sensing system in *S. pneumoniae* is activated in the murine respiratory tract, addition of activating CSP peptide pheromone was unable to induce competence in vivo [38].

In order to test the hypothesis that the Δ*rgg2* mutants were being killed before an inflammatory immune response was provoked, we examined bacterial counts and cytokine/chemokine quantification from the NALT (**Fig. 6**). Previous studies have shown that Il-1β and CXCL9 are upregulated in the NALT within twenty-four hours of intranasal infection with GAS [32, 33]. In a model of skin infection, colonization of wild type and MyD88-/- mice showed that the secretion of IL-1β and CXCL9 were dependent on MyD88 [34]. We found that IL-1β, CXCL9, as well as the proinflammatory cytokine TNF-α, were all downregulated in Δ*rgg2*-infected mice but not significantly different between mice infected with wild type or Δ*rgg3* (**Fig. 6D**). IL-1β has been shown to promote nasopharyngeal infection by GAS in the mouse model, and LaRock et al. suggest that this initial immune response and recruitment of neutrophils may be crucial for overcoming microbial interference [37]. The lack of activation of IL-1β in the *Δrgg2* bacteria lends credence to the hypothesis that QS-off bacteria are eliminated before eliciting MyD88-dependent activation of neutrophil migration to the NALT. As the Δ*rgg3* bacteria were seen to colonize the oropharynx more effectively than wild type, we expected that these mutants might induce cytokines and chemokines in the NALT to higher levels than seen for wild type. Curiously, the Δ*rgg3* bacteria elicited lower expression levels of several regulatory cytokines (IL-3, IL-4, IL-7) than wild type (**Fig. 6D**) even though mice were colonized with substantially higher CFU counts. Two of the Δ*rgg3*-infected mice had at least 40-fold more CFU in the NALT than mice colonized by wild type, yet cytokine levels overall were lower, suggesting that the Δ*rgg3* bacteria were affecting immune responses differently from wild type.

The inoculation of mice with mixed cultures of Δ*rgg2* and Δ*rgg3* bacteria intended to examine the consequences to GAS colonization when a portion of population (Δ*rgg2*) is unable to respond to pheromones produced by co-inoculated QS-active bacteria. Perplexingly, results of the mixed-culture inoculation experiments also showed that Δ*rgg3* mutants, which colonized to high levels in the pharynx, provided no benefit to Δ*rgg2* colonization (**Fig 5A**). These findings lead us to question how QS, which conceptually evolves to exert behaviors of cooperativity among community members, was in this case only benefiting GAS participating in QS. The exclusion of Δ*rgg2* mutants is reminiscent of QS policing mechanisms seen in *Pseudomonas aeruginosa*, where non-cooperative bacteria, or cheats, that exploit benefits shared by the community would incur a threat to the population if left unchecked [39, 40]. QS regulons that express “private good” genes, such as niche-specific catabolic pathways or stress response systems, confer advantages to individual bacteria, but only those that comply with social activities; bacteria that do not respond to pheromones or autoinducers (cheaters) will not induce these private goods and will therefore be less fit. Identifying the underlying mechanism via which Δ*rgg2* bacteria are unable to establish colonization remains to be determined. Due to this finding, the identification of quorum-regulated gene products retained by individual bacteria should be prioritized as possible mechanisms.

Many Rgg QS systems exist in gram-positive bacteria and, to date, the systems that have been studied all consist of a single transcriptional activator and a single *shp* pheromone [41].The Rgg2/3 system is unique in that there are two transcriptional regulators that antagonistically regulate the same targets. The entire system, including both *rgg* regulators, both *shp* pheromones and the two target downstream operons, is conserved throughout all sequenced strains of GAS. This conservation implies that the presence of two *rgg*s and two *shp*s is not a case of mere genetic redundancy but rather is important to the regulatory system’s architecture. The data in this work confirms the importance of both regulators, showing that the deletion of either *rgg2* or *rgg3* causes in vivo phenotypic divergence from wild type GAS behavior. Wild type bacteria appear to finely tune the expression of this system by using both regulators and the products produced by the system make great potential targets for vaccine development.

## Methods

### Bacterial strains

*S. pyogenes* strains used in this study and their method of construction can be found in Supplementary Table 1. *Streptococcus pyogenes* NZ131, MGAS315, and HSC5 were routinely cultivated in a chemically defined medium, CDM, as described previously [19] at 37^°^C and 5% CO2, without shaking. In order to facilitate timing of the experiments, starter cultures were prepared. Strains were grown overnight in THY broth, and in the morning, cultures were diluted 1:100 into CDM and grown to mid-logarithmic phase (OD600=0.5 to 0.6). Glycerol was added to a final concentration of 20% and aliquots of this culture were frozen and stored at −80° C. On experiment days, aliquots were diluted into fresh CDM 1:20 and grown to densities as specified below. In the study where bacteria were pretreated with synthetic peptide to activate the QS system before inoculation into the mouse, 100 nM SHP-C8 pheromone was added to the culture one hour before collection for inoculation.

### Construction of bacterial strains

The HSC5 Δ*rgg2* and Δ*rgg3* strains were constructed using the methods previously utilized to make the mutants in NZ131, as was the MGAS315 Δ*rgg2* strain [19].To delete the *rgg3* gene *(spyM3_0346)* in MGAS315, a 2,946 bp region encompassing *spyM3_0346* was amplified from MGAS315 genomic DNA using primers JC149 and JC155 and cloned into pFED760. Inverse PCR (primers JC309/JC310) followed by digestion with PacI allowed replacement of *rgg2* with an *aphA3* kanamycin resistance cassette and construction of the pJC244 plasmid (primers JC320/JC321; [42]).

### Murine colonization model

Female CD1 mice (Harlan/Envigo Laboratories) were obtained at age 3 weeks old. GAS bacteria were grown to an OD600 of 0.1 in CDM before centrifugation and resuspension in PBS. The mice were anesthetized with 3.5% isofluorane before intranasal inoculation in a 10 μl volume. The inoculum for MGAS315 and HSC5 was 1×10^7^ CFU while NZ131 was inoculated at 1×10^9^ CFU. At specified time points post-inoculation, the mice were again anesthetized with isofluorane and a calcium alginate swab (Calgiswab) was dampened with PBS and inserted into the oral cavity until resistance was met at the back of the throat. The swab was twirled in the oral cavity 10 times, and bacteria were subsequently released into PBS by vortexing the swab for 30 seconds. The resulting bacterial suspension was serially diluted and plated on BD Group A Selective Strep Agar with 5% Sheep Blood (BD Diagnostics) or THY agar plates with appropriate antibiotic selection, incubated at 37° C with 5% CO2 for 15 hours. Mice were monitored periodically by throat swabbing for up to 12 days post-infection. For NZ131 and HSC5, one trial of infections with 10 mice per bacterial strain is displayed. For MGAS315, two replicate trials of infections with 10 mice per bacterial strain were combined.

### IVIS Imaging of colonized mice

Infected mice were anesthetized using isofluorane and 1 μl of decyl aldehyde was instilled into each nostril before placement in a BSL-2 chamber within the IVIS instrument (IVIS Spectrum, Perkin Elmer). Isofluorane anesthesia was sustained during IVIS imaging. Biosafety regulations limited imaging to be taken no earlier than 15 minutes post-inoculation; a second measurement was taken at 60 minutes. The mice were released into a cage unanesthetized between the two timepoints. Standard operating procedure approved by UIC was used to maintain a BSL-2 environment during the course of the experiment.

### Quantitative PCR from broth culture

Starter cultures were diluted into CDM and grown to an OD of 0.1, after which 100nM SHP-C8 or 100nM REV-C8 were added. Cells were harvested between OD 0.3-0.4 and RNA was isolated and purified using the Ribopure Bacteria Kit (Invitrogen). Samples were treated with DNase I to eliminate genomic DNA. Purified RNA was converted to cDNA using the Superscript III First-Strand Synthesis System with random hexamers as the amplifying primer (Thermo Fisher Scientific). qRT-PCR was done using the Fast SYBR Green Master Mix (Applied Biosystems) and a CFX Connect Real Time PCR detection system (Bio-Rad). Control reactions omitting the reverse transcriptase were also performed to eliminate the possibility that signal was due to contaminating genomic DNA. The gyrase A gene (*gyrA*) was utilized as a housekeeping gene [43–45]. Samples were run in biological triplicate, and all samples were run in technical triplicate on a single plate. Statistical significance of each condition compared to wild type grown in broth was determined by using the Holm-Sidak method, with α=0.05.

### Quantitative PCR from the mouse

Mice were anesthetized with isofluorane before inoculation with 1×10^8^ CFU of MGAS315. One-hour post-inoculation, the mice were again anesthetized with isofluorane, and a calcium alginate swab was inserted into the oral cavity and twirled 10 times. The swabs were then incubated in 1 ml Trizol at room temperature, vortexed for one minute and sonicated for three minutes before being subjected to RNA purification via the Ribopure Bacteria Kit (Invitrogen). Swabs from 10 mice were pooled in order to isolate sufficient amounts of RNA, which has been cited as a stronger sampling strategy when compared to unpooled samples with a low limit of detection [46]. Cells growing in CDM with and without pheromone were subjected to RNA purification to act as controls for the mouse samples. Samples were subsequently treated with DNase I to eliminate genomic DNA. Purified RNA was converted to cDNA using the Superscript III First-Strand Synthesis System (Thermo Fisher Scientific), including treatment with RNase H. Antisense gene-specific primers KMT044 (*gyrA*), KL1 (*stcA*), and 450R (*aroE.2*) were used for cDNA synthesis; sequences can be found in Supplementary Table 4. qRT-PCR was done using the Fast SYBR Green Master Mix (Applied Biosystems) and a CFX Connect Real Time PCR detection system (Bio-Rad). Control reactions omitting the reverse transcriptase were also performed to eliminate the possibility that signal was due to contaminating genomic DNA. The gyrase A gene (*gyrA*) was utilized as a housekeeping gene [43–45]. Samples grown in culture were run in biological triplicate; as mentioned above, in vivo samples were prepared from 10 mice pooled to retrieve useable levels of RNA. All samples were run in technical triplicate on a single plate, and statistical significance of each condition compared to wild type grown in broth was determined by using the Holm-Sidak method, with α=0.05.

### NALT bacterial counts and multiplex cytokine array

10 mice were infected with MGAS315 WT, Δ*rgg2,* or Δ*rgg3* as previously described. Five mice from each condition were subjected to euthanasia and NALT surgery 24 hours post-infection; the other five were processed 72 hours post-infection. The NALT organs were removed as previously described [47]. One half of the bi-lobed organ was homogenized and plated to enumerate bacterial counts. The other half was processed to extract supernatants by mashing the tissue against a 24-well plate with the back of a syringe. The resulting mixture was resuspended in 150 μl of sterile PBS. The supernatant was then clarified by centrifuging the suspension at 17000× *g* for 10 minutes at 4° C. 5 μl of the supernatant was used to normalize the level of total protein in each sample via BCA assay (BioRad) and the supernatant was then flash frozen on dry ice. Supernatants were submitted to the UIC Flow Cytometry Core in order to carry out a 32-plex cytokine/chemokine analysis using the 32-plex Milliplex Map Mouse Cytokine/Chemokine Magnetic Bead Panel (Millipore). Results were compared between WT and Δ*rgg2* or WT and Δ*rgg3* by applying a student’s t test (p=0.05).

### Ethics Statement

The mice and methods used in this work were approved by the Institutional Animal Care and Use Committee (IACUC) at UIC, which is known as the Animal Care Committee (ACC). The ACC is responsible for the review and approval of all research, testing and teaching using animals at UIC. The campus recognizes the following regulatory authorities for the care and use of animals: Association for Assessment and Accreditation of Laboratory Animal Care - International (AAALAC), the U.S. Department of Agriculture (USDA), and the Office for Laboratory Animal Welfare (OLAW). The protocol writing and review process is administrated through the Office of Animal Care and Institutional Biosafety (OACIB) within the Office of the Vice Chancellor for Research. OACIB reviewed and accepted the protocols utilized in this study, under the identification number of 17-013.

## Acknowledgments

Work herein was supported by funds associated with NIH R01-AI091779. AG is supported by F30AI136359 and MF by the Burroughs Wellcome Fund for Investigators of Pathogenesis of Infectious Diseases. We thank Dr. Jennifer Chang for previous construction of the Δ*rgg2* and Δ*rgg3* mutant strains in the MGAS315 and HSC5 backgrounds. They were invaluable to this study. Thank you to Kayleigh Tovar and Kaitlyn Tovar for assistance during the IVIS experiments.

## Supporting information captions

**Figure S1: Colonization of the murine oropharynx by S. pyogenes strains NZ131 and HSC5**

**Table S1: Raw Numbers from Milliplex Cytokine Array from Day 3 (pg/ml).** Data correspond with Figure 6AB of the main text.

**Table S2: Bacterial Strains Used in this Study.**

**Table S3: Plasmids Used in this Study.**

**Table S4: Primers Used in this Study.**

## Supplemental Information

**Figure S1.**
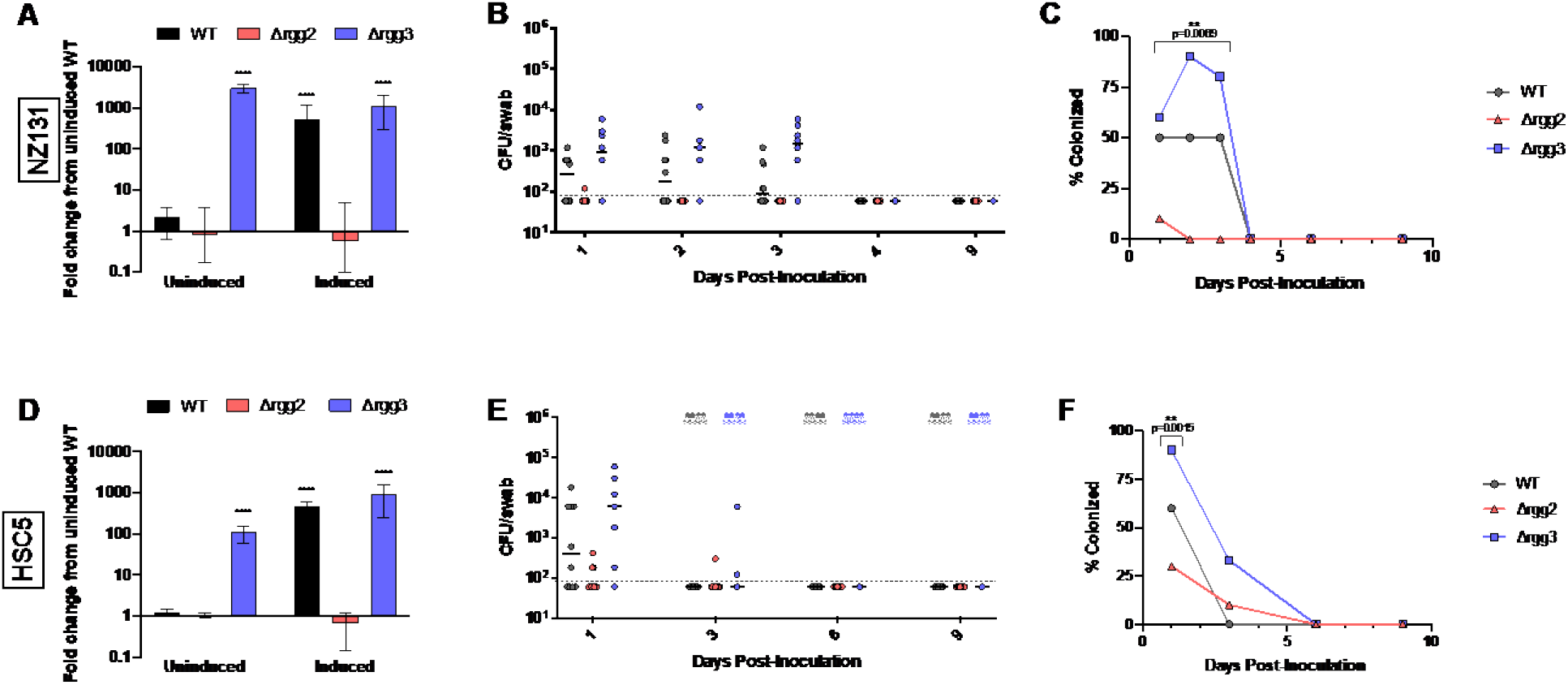
Colonization of the murine oropharynx by *S. pyogenes* strains NZ131 and HSC5. GAS strains NZ131 and HSC5 were analyzed for QS activity in culture and for colonization levels in the mouse. (A/D) Relative transcript levels of QS-target gene *stcA* in cultures of WT, Δ*rgg2*, and Δ*rgg3* strains grown in CDM broth without (uninduced) or with (induced) the addition of 100 nM SHP-C8 pheromone. Bars are the mean fold differences compared to uninduced WT for three biological replicates. Asterisks indicate significance of multiple t-tests comparing each sample to uninduced WT, α≤0.05. (B/E) Three-week old CD1 mice were inoculated intranasally with 10^9^ CFU of uninduced bacteria. CFU counts obtained from throat swabs of each mouse are plotted for each day of sampling. Skull and crossbones denote the number of animals that displayed humane endpoint signatures requiring euthanasia (signs of distress indicating invasive infection) and are shown in an accumulating manner. Ten mice were utilized for each bacterial strain; bars indicate the median CFU/swab value and the detection limit was 60 CFU/swab. Using the Friedman test, a nonparametric matched ANOVA, the NZ131 strains were determined to have statistically different median CFU counts over the first three days of the experiment, p=0.028. The HSC5 strains were compared for the first day using the Friedman test and had statistically significant median CFU counts, p=0.0015. (C/F) The percentage of mice colonized during the experiment shown in panels B and E were compared between the three strains over time. Deceased mice were not included during the calculation of percent colonized mice. A Friedman test was again run to compare the colonization percentages between NZ131 three strains on the first three days, and the colonization rates of each strain were found to be statistically significant from one another, p=0.0069. The first day’s percentage of colonized HSC5 strains were also compared using a Friedman test, no significance was found (α≤0.05).

**SUPPLEMENTAL TABLE 1.**
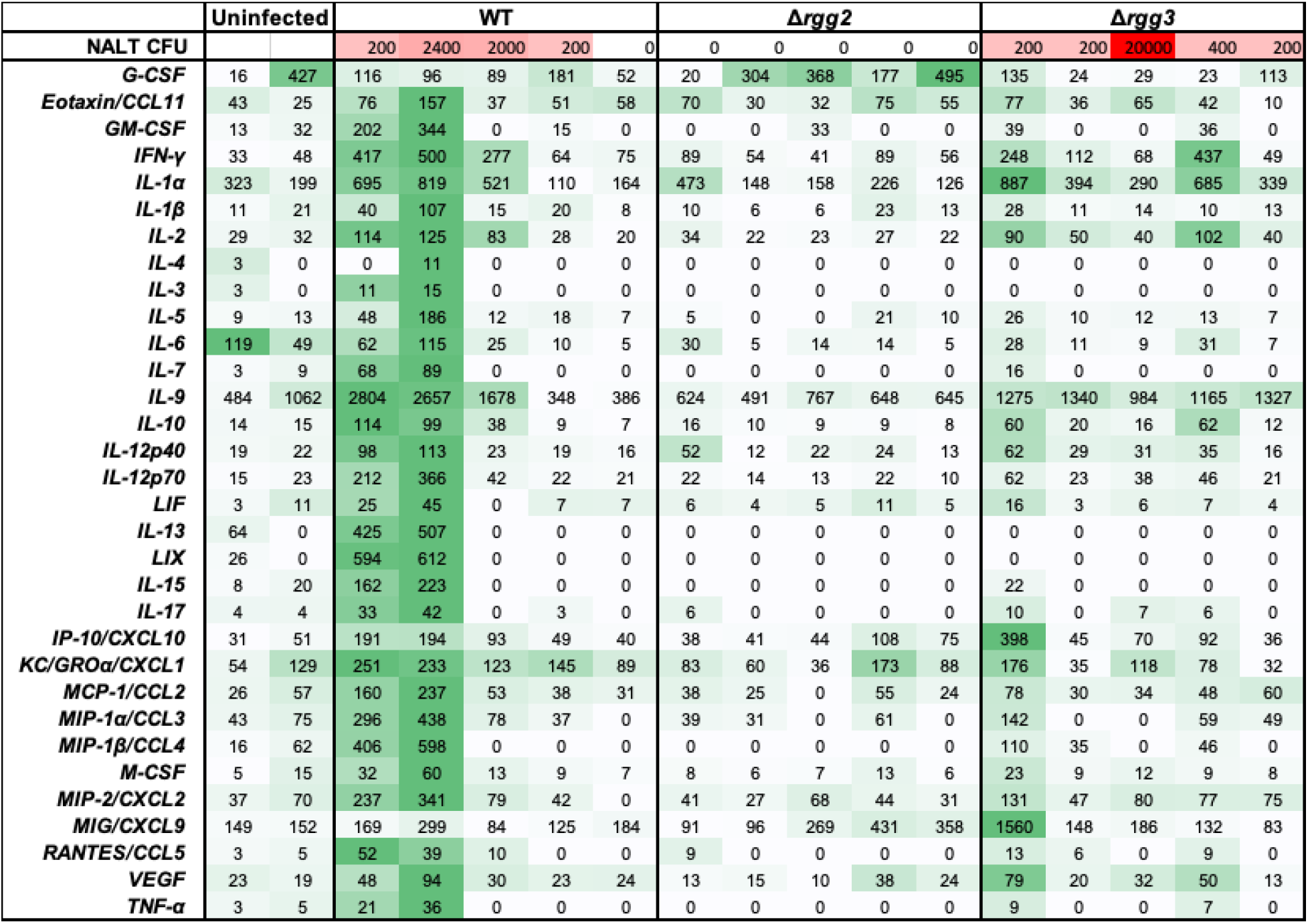
Milliplex Assay Raw Numbers from Day 3 (pg/ml). Data correspond with Figure 6AB of the main text.

**SUPPLEMENTAL TABLE 2.** Bacterial Strains Used in this Study.

| Strain Name | Description | Reference |
| --- | --- | --- |
| NZ131 | *S. pyogenes* M49 serotype | [1] |
| MGAS315 | *S. pyogenes* M3 serotype | [2] |
| HSC5 | *S. pyogenes* M14 serotype | [3] |
| JCC137 | NZ131 $\Delta rgg2$; unmarked | [4] |
| JCC131 | NZ131 $\Delta rgg3::cat$; CmR | [4] |
| JCC196 | MGAS315 $\Delta rgg2::aphA3$; KanR | This study |
| JCC194 | MGAS315 $\Delta rgg3::cat$; CmR | This study |
| JCC195 | HSC5 $\Delta rgg2$; unmarked | This study |
| JCC190 | HSC5 $\Delta rgg3::cat$; CmR | This study |

**SUPPLEMENTAL TABLE 3.** Plasmids Used in this Study.

| Plasmid | Description | Backbone | Source |
| --- | --- | --- | --- |
| pJC162 | Temperature sensitive backbone, ErmR | pFED760 | [4] |
| pJC175 | to replace *rgg3* with cat cassette in NZ131/HSC5 | pJC162 | [4] |
| pLA101 | *rgg3* region cloned into pJC162 | pJC162 | [4] |
| pOsKar | Source of the kanamycin cassette |  | [5] |
| pJC243 | MGAS315 *rgg2* region cloned into pJC162 | pJC162 | This study |
| pJC244-Kan | *Kan* cassette replacing *rgg2* gene in pJC243 | pJC243 | This study |
| pJC229 | P<sub>shp3</sub>-*luxAB* luciferase reporter on multicopy plasmid | pLZ12-Sp | [6] |

Cm: Chloramphenicol, Kan: Kanamycin, Sp: Spectinomycin, Erm; Erythromycin

**SUPPLEMENTAL TABLE 4.**
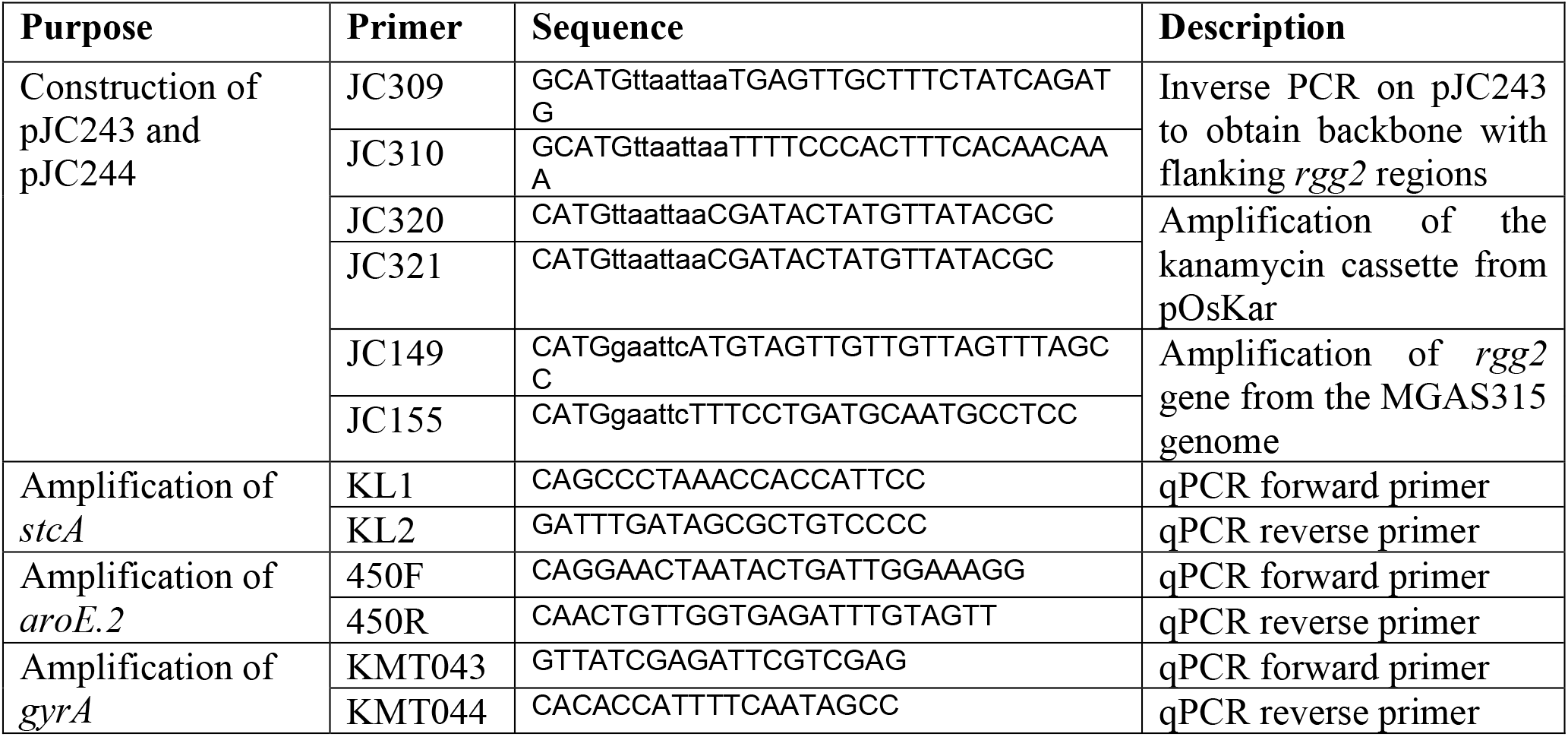
Primers Used in this Study.

